# Biology Undergraduates’ Use of AI-Enabled Feedback to Revise Scientific Writing: Recipience and Barriers

**DOI:** 10.64898/2026.10.19.760216

**Authors:** Michele Weston, Tammy Long

## Abstract

**Background:** Instructors and education researchers are increasingly developing feedback with the assistance of generative AI, which we refer to as AI-enabled feedback, yet we do not know whether established understandings of feedback uptake and effectiveness apply similarly in AI-mediated contexts. This makes the recipience of AI-enabled feedback a central concern to understanding how to best support students in developing their scientific writing skills. This study examined how two teams of undergraduate biology students used AI-enabled feedback to revise their scientific writing, and the extent to which their revisions addressed that feedback. The teams revised their research proposal drafts after receiving AI-enabled feedback developed by their instructor with the aid of a custom AI tool.

**Results:** Our findings show that AI-enabled feedback prompted actionable revision across both teams. Each team addressed approximately half of the feedback comments, indicating comparable levels of uptake despite their differing approaches to revision. We observed that both teams were inconsistent in providing the additional information and reasoning requested in the feedback, and they struggled to address comments requiring self-evaluation. Taken together, we see that students often addressed the more easily actionable parts of AI-enabled feedback while omitting more demanding revisions.

**Conclusions:** These findings provide behavioral evidence of how students used AI-enabled feedback to revise their scientific writing. The observed patterns suggest that the content of AI-enabled feedback supported meaningful revision, while limitations in students’ agency, strategies, and volition likely shaped their implementation of comments. Although the degree of uptake observed in this study is encouraging, it is unlikely that a single application of generative AI within the feedback process will resolve all the challenges associated with feedback uptake.

---

Formative feedback on learning tasks is understood to be one of the most influential factors contributing to student achievement (Black & Wiliam, 1998; Hattie, 1999). In undergraduate science education, feedback plays an important role in helping students develop scientific writing, a complex practice that requires students to apply disciplinary conventions while communicating for a specific purpose and a scientific audience (Rivard, 1994). For feedback to be most effective in supporting students’ learning, they must actively engage with it and use it to revise their work (Boud & Molloy, 2013). This makes feedback uptake and the factors influencing it a central concern to understanding how to best support students in developing their scientific writing skills. As generative artificial intelligence (genAI) is increasingly used to provide student-facing feedback on writing tasks, it is becoming important to understand how these technologies shape students’ engagement with feedback during the writing process. This perspective shifts emphasis away from evaluating the capabilities of particular genAI tools and toward understanding how this feedback influences students’ learning processes and writing outcomes.

## Barriers To Feedback Use

To explain why feedback is more effective in some contexts than others, researchers have increasingly focused on barriers to feedback use. Feedback can be regarded as a process that occurs in three phases: instructors’ capacity to provide feedback, students’ understanding of feedback, and students’ implementation of feedback. Figure 1 illustrates our categorization of barriers according to when they occur in the feedback process.

**Figure 1.**
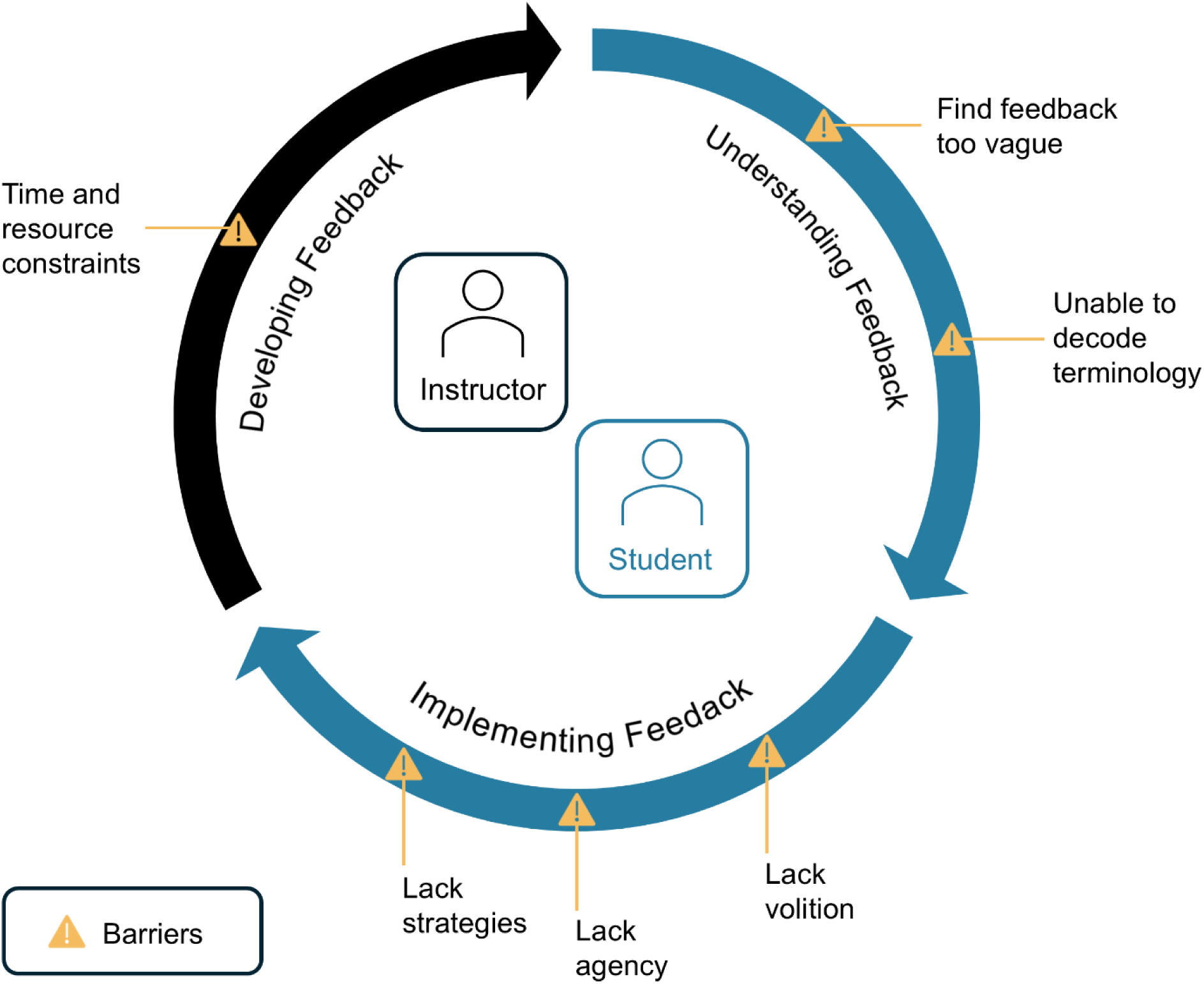
Learner-centered feedback process (Boud & Molloy, 2013; Henderson et al., 2019) and barriers to feedback use.

Barriers related to instructor capacity arise when students receive little feedback or encounter delays in receiving it. In some cases, students receive little information about their performance, or no feedback at all (Derham et al., 2022; Jørgensen et al., 2023). Even when feedback is provided, delays in its delivery can reduce students’ engagement with it (Blair et al., 2013; Nicol & Macfarlane-Dick, 2006; Yang & Carless, 2013). These barriers are often because instructors face time and workload constraints that limit their ability to provide timely, personalized feedback (Heim & Holt, 2019).

Barriers to feedback use may also emerge after students receive feedback, and often are related to students’ ability to understand and interpret feedback. Students frequently report difficulty understanding feedback, particularly when comments are vague or contain unfamiliar terminology or symbols. When feedback comments aren’t specific or detailed enough, such as short statements or the use of symbols and underlining, students report not having enough information to know what the error is or what to work on (Jørgensen et al., 2023; Sutton, 2012). In contrast, students report that personalized and detailed feedback supports comprehension by making expectations and areas for improvement more explicit (Dawson et al., 2019). Understanding may also be hindered by limited familiarity with academic or discipline-specific terminology, whether those terms appear in feedback comments themselves or in the evaluative criteria to which the feedback refers (Jonsson, 2013; Sutton, 2012).

However, comprehension alone does not ensure effective feedback use. Even when students understand feedback, they may struggle to translate comments into next steps that will improve their work (Winstone, Nash, Rowntree, et al., 2017). Research in this area has identified several factors that contribute to this difficulty. Most notably, some students lack strategies for implementing feedback, leading them to engage with comments passively or remove problematic sections of their work rather than revise them (Hyland, 1998; Winstone, Nash, Rowntree, et al., 2017). Others may be unwilling to invest the effort required to make substantive improvements (Sutton, 2012; Winstone, Nash, Parker, et al., 2017). Without motivation for pursuing improvement and development as a learner, students are unlikely to engage with feedback in ways that support its implementation (Winstone, Nash, Rowntree, et al., 2017).

## Learner-Centered Feedback Process

Conceptualizations of feedback have evolved over time. Traditionally, feedback has been viewed as information about task performance (e.g., written evaluative comments) that is provided to students (Hattie & Timperley, 2007). This transmission-oriented view treats feedback as largely unidirectional and pays limited attention to what happens after students receive feedback. Consequently, research adopting this perspective tends to focus on the content of feedback and its delivery, such as timing and mode. This part of the feedback process is shown as ‘Developing Feedback’ in Figure 1. In contrast, modern perspectives conceptualize feedback as an active, learner-centered process in which students use information about their performance on a task to promote their learning (Boud & Molloy, 2013; Henderson et al., 2019). From this perspective, feedback is effective not because information is provided, but because learners are able to understand, interpret, and act on that information. As a result, researchers have increasingly focused on students’ experiences with feedback, examining how they perceive, engage with, and use feedback to support learning (e.g., Dawson et al., 2019; Jonsson, 2013; Li & De Luca, 2014). Figure 1 presents these additional components of the process as ‘Understanding Feedback’ and ‘Implementing Feedback’. We adopt this learner-centered perspective in the present study. From this perspective, the effectiveness of feedback depends on how learners engage with and use feedback.

Winstone, Nash, and Rowntree et al. (2017) advanced a learner-centered view of feedback through the concept of proactive recipience, which is when a student takes an active role in engaging with feedback. Proactive recipience shapes how students experience and respond to feedback throughout the feedback process, while features of the feedback and its context can, in turn, influence students’ level of recipience (Winstone, Nash, Parker, et al., 2017; Winstone, Nash, Rowntree, et al., 2017). Studies that informed this view documented differences in how students engage with feedback. In some cases, students do not read or use feedback (Bevan et al., 2008; Cohen, 1987; Furnborough & Truman, 2009; Hyland, 1998), whereas in others they incorporate feedback into subsequent work (Higgins et al., 2002; Winstone, Nash, Parker, et al., 2017). In the context of writing, acting on feedback often involves revising a draft. Yet revisions do not always result in meaningful improvement. Students often focus on surface-level changes, such as spelling, grammar, or minor details, rather than more substantive improvements to the quality of the work (Ashwell, 2000; Zhu et al., 2020). Such variation in students’ use of feedback suggests that factors influencing recipience may play an important role in determining the effectiveness of feedback.

## Generative AI Use in Feedback Development

Instructor feedback plays a crucial role in supporting students’ scientific writing, particularly when there are opportunities to reflect and revise (Jerde & Taper, 2004; Yeh, 2010). GenAI is emerging as a promising tool for assisting instructors in developing formative feedback on students’ scientific writing, such as lab reports (Mills et al., 2025; Wan & Chen, 2024), and shorter constructed-response tasks (Chu et al., 2026; Cooper & Klymkowsky, 2024; Pellegrino, 2024). GenAI refers to artificial intelligence systems capable of producing original content, such as written text, in response to user prompting (Stryker & Scapicchio, 2024). Compared to traditional machine learning (ML) approaches, genAI systems offer advantages for feedback generation: its natural-language prompting makes it easy to interact with, and its pretrained large language models allow it to perform tasks without requiring the large, scored datasets typically needed for supervised ML systems. These affordances create a low barrier to entry and position genAI to be adopted more broadly by instructors.

Given the growing use of genAI to support feedback development, recent studies have examined students’ perceptions of the feedback it helps produce. Findings have been mixed. In some contexts, students find AI-supported feedback on lab reports and project proposals to be useful, actionable, and clearly written (Mills et al., 2025). In other cases, however, students perceive the feedback less favorably. For example, Lim and Go (2025) reported that students found AI-generated feedback on essays to be overly broad, generic, and vague. One possible explanation for these differences lies in how AI systems are designed and implemented. Because Lim and Go relied on a standard ChatGPT prompt without additional instructions or a domain-specific knowledge base, the resulting feedback largely consisted of generalized comments resembling grading criteria. In contrast, Mills et al. (2025) configured a customized version of ChatGPT that had specialized instructions and access to a curated knowledge base, thus generating detailed and consistent feedback. These findings suggest that variation in students’ perceptions may reflect variation in AI tools and approaches.

### Promoting Quality AI-Enabled Feedback

As genAI becomes more widely applied in educational settings, it is important to foster ethical and effective use by intentionally aligning AI implementations with instructional goals and values. Human-centered AI offers one such approach by emphasizing design practices that augment rather than replace human capabilities and that maintain human oversight of AI systems (Schmager et al., 2025). We define AI-enabled feedback as feedback developed within a human-centered AI process. Keeping instructors actively involved in the feedback process helps ensure higher-quality output and preserves accountability for assessment. In practice, human involvement may include configuring and testing an assignment-specific AI system, as well as reviewing and refining AI output before sharing it with students. Customizing genAI systems involves providing the instructions and resources to a model to perform a specific task that reflects the course context, learning goals, and assessment criteria. Such customization enhances the accuracy of feedback and reduces hallucinations, which are instances of the model producing inaccurate yet plausible text (Mills et al., 2025; OpenAI et al., 2024). Viewed in this way, the value of AI-enabled feedback lies not in automating the process, but in supporting instructors’ ability to provide quality, personalized feedback to all students.

## Research Goals

GenAI tools are increasingly being used by instructors to automate, augment, and expand the feedback they provide to students (e.g., Kasneci et al., 2023; Mills et al., 2025; Wan & Chen, 2024; Xia et al., 2026). However, we do not yet know whether established understandings of feedback uptake and effectiveness apply in AI-mediated contexts (Cooper, 2026; Kasneci et al., 2023). Understanding students’ recipience of AI-enabled feedback is therefore a crucial first step toward evaluating its role in supporting scientific writing. In this study, we investigate biology undergraduates’ recipience of AI-enabled feedback during the revision of scientific writing. To do so, we examine how students used AI-enabled feedback and the extent to which they addressed that feedback in their revisions. Our research questions are:

1) How did students use AI-enabled feedback to revise their scientific writing?
2) To what extent did students address the AI-enabled feedback?

## Methods

To investigate students’ recipience of AI-enabled feedback, we used a qualitative, multiple case study design and analyzed students’ revisions to their scientific writing. Multiple case studies examine a phenomenon within bounded contexts, or cases, in order to focus on the experiences of participants (Merriam, 1998; Miles & Huberman, 1994). This approach emphasizes an in-depth analysis of a small number of cases, making it ideal for presenting insights that inform our understanding of the nuance of students’ use of AI-enabled feedback. The study includes two cases, each consisting of a team of three students.

### Study Context

This study was conducted within a STEM-focused residential college at a large research university located in the Midwest. The research took place in the laboratory component of a combined lecture and laboratory course on introductory organismal biology. The laboratory curriculum was designed to explore systemic inequities and human impacts on ecology and health. The first part of the course introduced students to different ecological systems, methods, and types of data analysis. Following this, student teams developed independent research projects investigating the underlying causes of a human health condition of their choice and its relationship to an ecological disparity. Although research projects often utilize final papers resembling journal articles, in this course, each team was required to incorporate their results into a written research proposal as preliminary findings that will inform a new project. While the original intention was for students to complete full research proposals, time constraints required the assignment to be shortened to the introduction section only. Therefore, we refer to the assigned piece of writing as the “proposal introduction”. Students were instructed to follow a rubric that specified the content of four required paragraphs, each focusing on a different scale of human impact. Descriptions of the paragraphs are shown in Table 1.

**Table 1.**
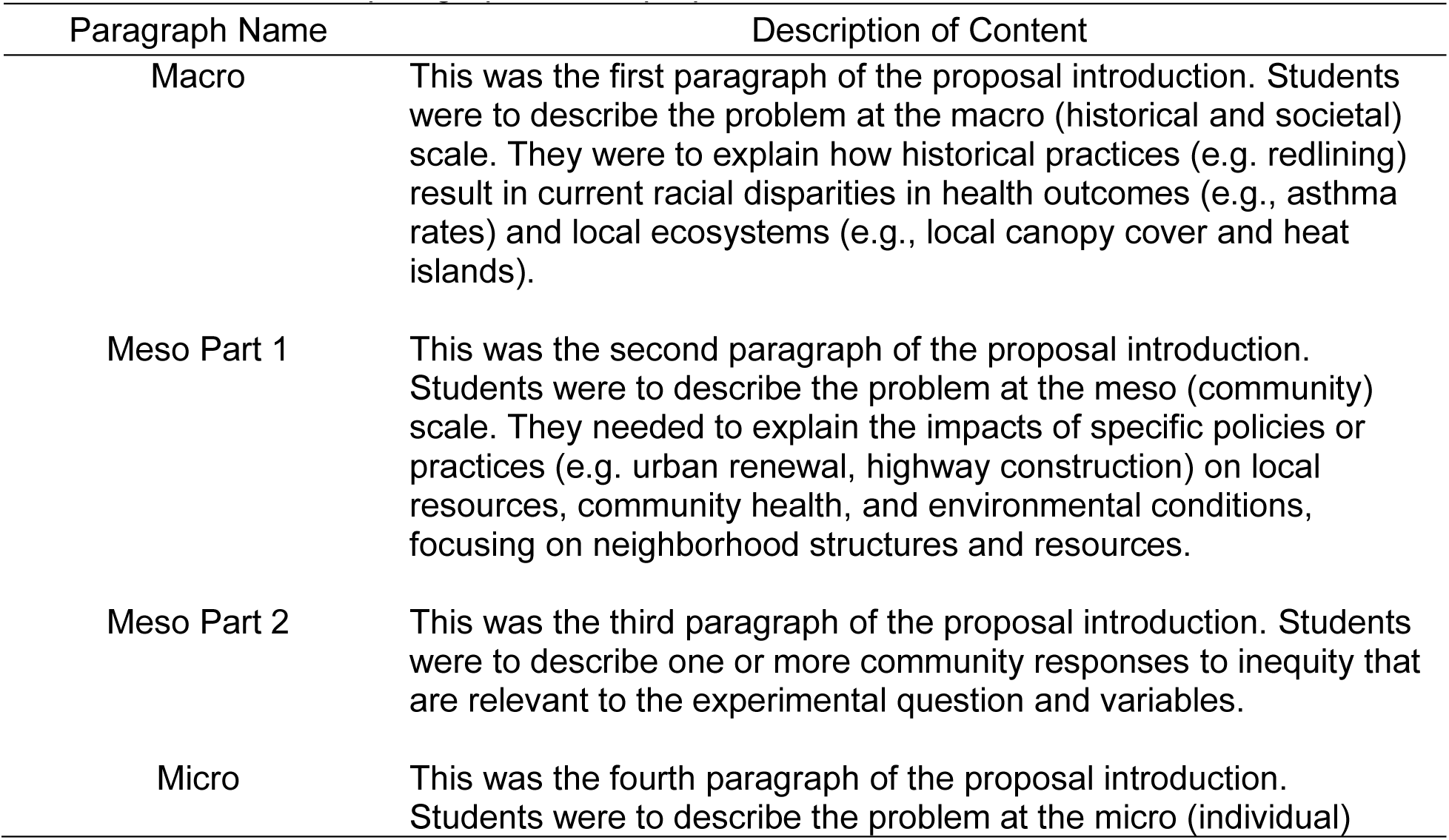
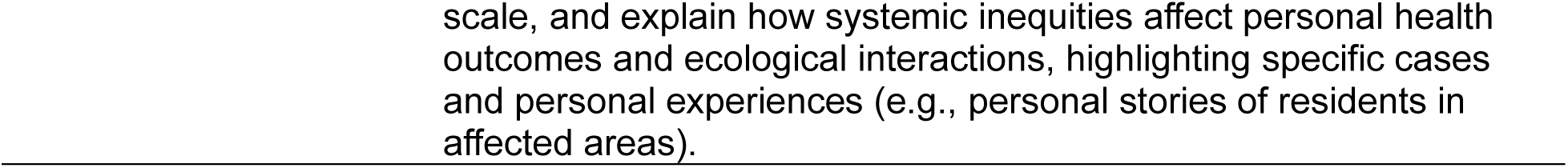
Content of the paragraphs of the proposal introduction.

### Participants

All students enrolled in the laboratory course were invited to participate in the research study. There were two student teams where all members consented to participate, and they were each considered a case. Table 2 provides demographic and academic information for these participants.

**Table 2.** Student participant demographics.

| Participant | Team Number | Gender | Race(s) and Ethnicity/ies | Final Course Grade <sup>a</sup> | Cumulative GPA <sup>b</sup> |
| --- | --- | --- | --- | --- | --- |
| Student 1 | 1 | Woman | White | 4.0 | 3.83 |
| Student 2 | 1 | Man | White | 3.0 | 2.75 |
| Student 3 | 1 | Woman | South Asian | 4.0 | 3.79 |
| Student 4 | 2 | Man | Black or African American | 4.0 | 3.59 |
| Student 5 | 2 | Woman | Black or African American | 4.0 | 2.44 |
| Student 6 | 2 | Woman | White | 4.0 | 4.0 |

<sup>a</sup> Final course grade is reported on a 4.0 scale. <sup>b</sup> Cumulative GPA is reported on a 4.0 scale and at the start of the semester in which the study took place.

### Procedures

Student teams began the assignment by drafting the proposal introduction and submitting it to their instructor. The instructor then used a genAI tool (described below) to develop customized feedback aligned with the assignment rubric, reviewed the feedback for accuracy and quality, and made edits as necessary before providing it to students. The instructor had the option of using follow-up prompts to refine the genAI tool output but opted not to do so. Students then submitted the revised proposal introduction as the final draft.

### Generative AI Tool

This study utilized a GPT, which is a specialized version of ChatGPT designed to carry out a specific task (*Introducing GPTs*, 2024). Although it resembled the standard ChatGPT interface, allowing the instructor to interact with a chatbot through a chat window, the GPT used in this study (model 4o; OpenAI, 2025) was configured with detailed instructions and a curated knowledge base. As a result, it could query course-relevant documents and incorporate those materials into its responses. This additional context can reduce hallucinations by grounding the model’s predictions in relevant information (*Introducing GPTs*, 2024; *What Is RAG?*, n.d.). Unlike prompts entered into a standard chat session, GPTs retain their instructions and reference materials across sessions, allowing users to repeatedly perform the same specialized task without re-entering those materials. Additional details about the GPT are provided in the Supplemental Materials.

The GPT was intentionally grounded in learning theory and course-specific supports. The lab coordinator of the course and the first author collaborated to design the research proposal assignment and its rubric, ensuring alignment with the coordinator’s pedagogical priorities. The rubric incorporated the Claim, Evidence, and Reasoning (CER) framework, which had been used throughout the course to support students’ scientific explanations (McNeill & Krajcik, 2008). To further guide students, we developed a scaffolding tool that outlined the structure of the proposal introduction and specified the content expected in each paragraph. Building on this foundation, we configured and tested the GPT to provide feedback tailored to the rubric, CER framework, and scaffolding tool. As a result, the feedback mirrored the organization of the rubric, with the structure described in Table 3. The resulting feedback was structured so that each rubric section included (1) identified strengths, (2) suggestions for improvement, and (3) examples intended to scaffold those improvements. Student data was protected by collecting written consent for the use of genAI in analyzing student work, anonymizing proposals prior to use of the GPT, and ensuring that proposals were not retained or used by OpenAI to train or improve its models.

**Table 3.** Structure of the feedback sections provided to students.

| Feedback Section | Scope | Description |
| --- | --- | --- |
| Macro | Paragraph-specific | Feedback pertaining to the content of the Macro paragraph. |
| Meso | Paragraph-specific | Feedback pertaining to the content of both the Meso Part 1 and Meso Part 2 paragraphs. |
| Micro | Paragraph-specific | Feedback pertaining to the content of the Micro paragraph. |
| Flow of Writing | Whole-proposal introduction | Feedback on how effectively each sentence and paragraph transitions to the next, with a clear flow of ideas. |
| Claims are Supported | Whole-proposal introduction | Feedback on how well claims are supported by scientific reasoning and evidence from credible cited sources. |

*Note.* Each section followed the standardized format of strengths, suggestions for improvement, and scaffolding tips.

### Data Analysis

Our revision analysis included two stages. First, we compared the first and final drafts to identify and categorize the types of changes. Then, we cross-referenced the AI-enabled feedback with the changes to examine how each team used their feedback. We adapted and applied a revision operations coding scheme (Faigley & Witte, 1981) to the drafts to categorize the meaning-based revisions as additions, deletions, moves, and substitutions. A change was considered meaning-based if it involved conceptual ideas or information. Additions and deletions of citations were coded separately to distinguish them from meaning-based revisions. Surface-level revisions were excluded from analyses and included changes that corrected spelling or grammar, or edits that influenced the implicit or explicit nature of an idea without changing the idea itself. When a revision was longer than a single sentence, we coded each sentence separately. One exception occurred with Team 1, whose first draft was missing one of the required paragraphs. Because there was no original text to revise, the corresponding paragraph in the final draft was not coded for revisions. This ensured that the revision analysis focused only on changes to existing content. Table 4 presents illustrative examples of the revision operations coding [Additional file 1]. The Supplemental Materials include both teams’ complete drafts annotated with our revision coding.

**Table 4.**
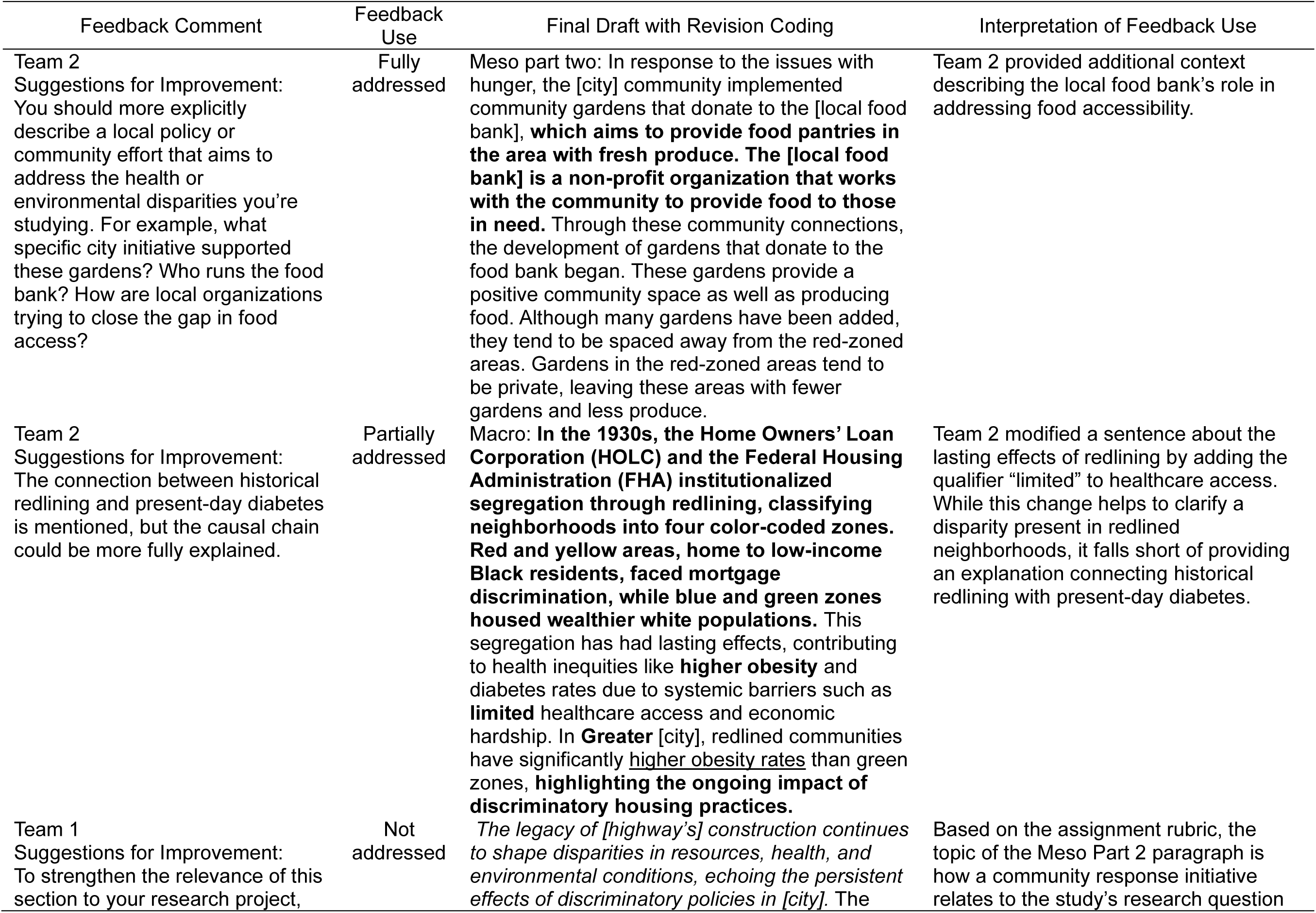
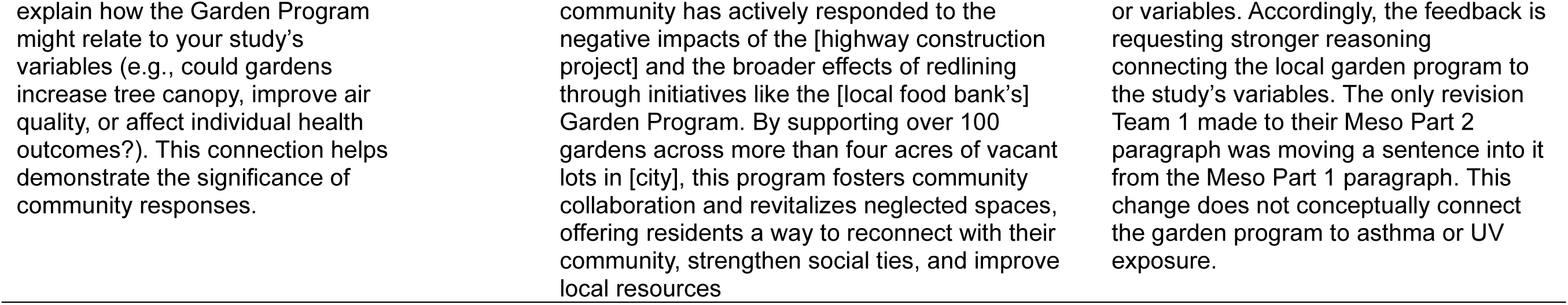
Illustrative examples of feedback use.

After categorizing the types of changes teams made, we then examined the uptake of each feedback comment. For each comment, we identified the relevant segments in both the first and final drafts and categorized the comment as fully addressed, partially addressed, or not addressed based on the extent to which the suggested changes were reflected in the final draft. A comment was categorized as partially addressed when the revisions showed progress toward addressing the issue but did not fully resolve it. This analysis focused only on feedback comments that offered a suggestion for improvement or a scaffolding tip (e.g., examples of how to improve). Comments highlighting strengths were excluded because they did not require revision. We conducted all coding and analysis in MaxQDA (VERBI Software, 2024). Table 4 provides examples of feedback comments, their uptake classification, and corresponding excerpts from student drafts.

## Results and Discussion

We begin this section with the results of the revision analysis including students’ use of feedback comments. That is followed by within-case analyses of each team to illustrate and contextualize the results of the revision analysis, and a cross-case analysis synthesizing the findings. We end the section with theoretical interpretations of the findings.

### Revision Analysis

Our revision analysis identified and categorized types of meaning-based revisions that students made after receiving AI-enabled feedback. Across both teams, all four types of meaning-based revisions were observed. Deletions were the most frequent (14 of 31 changes), followed by additions (10 of 31 changes). Team 1 made more revisions overall, even though one missing paragraph in their first draft meant that fewer paragraphs were included in the revision analysis. The teams also differed in their distribution of meaning-based revisions: Team 1 primarily deleted content, whereas Team 2 made more additions than deletions. Notably, both teams removed citations from their first drafts and did not introduce any new citations in their final drafts. Figure 2 presents a detailed breakdown of revision operations used by each team.

**Figure 2.**
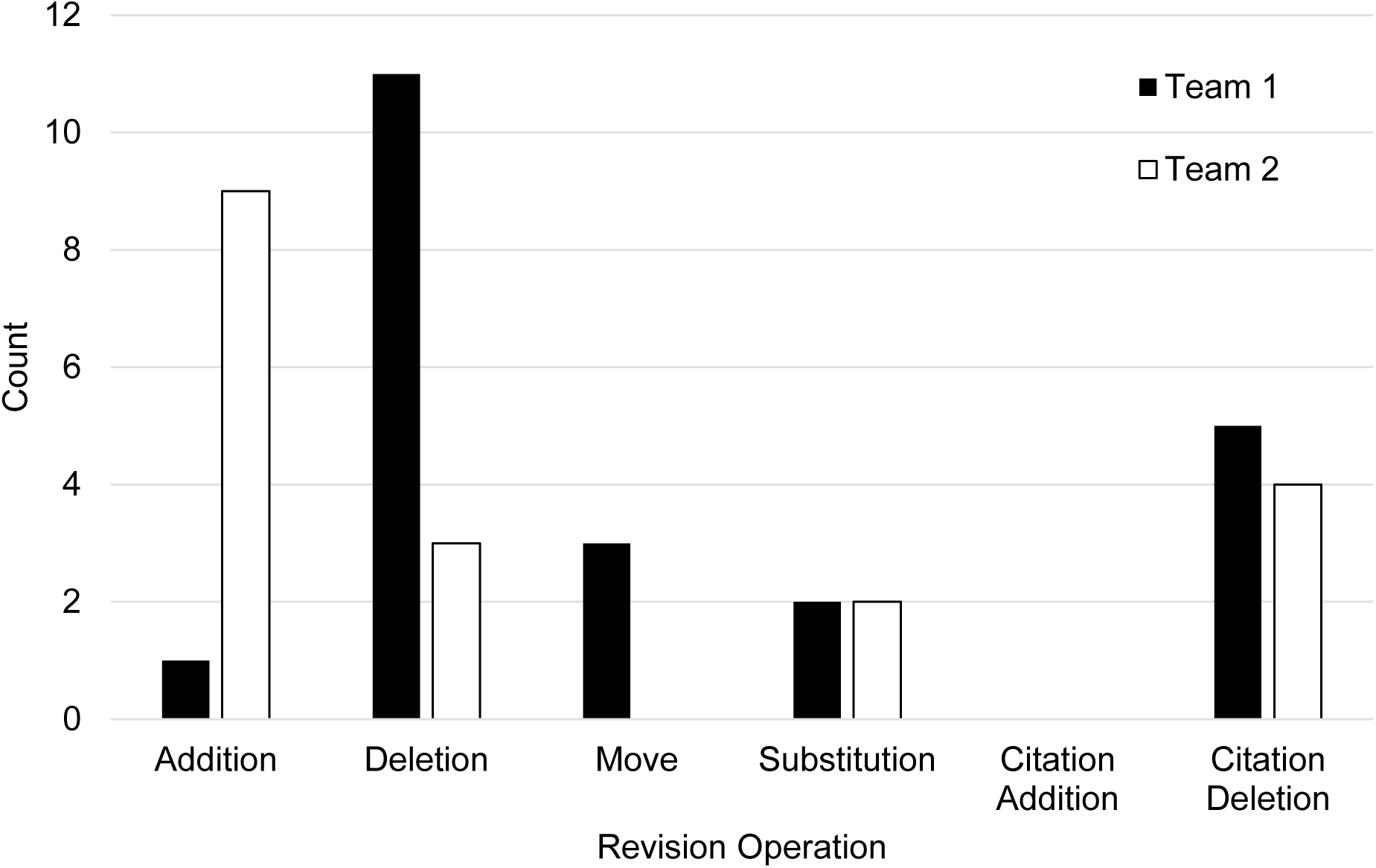
Count of revision operations for each student team.

To examine how student teams incorporated AI-enabled feedback into their revisions, we classified each feedback comment as fully, partially, or not addressed, based on the extent to which suggested changes were reflected in the revised draft. Because Team 1’s initial draft was missing a required paragraph, they had fewer feedback comments included in the analysis. Across both teams, 10 of the 22 comments were fully or partially addressed, leaving more than half unaddressed. The distribution of comments across the categories fully, partially, and not addressed was similar for both teams and consistent across comment type (suggestions for improvement vs. scaffolding tips). Figure 3 presents the distribution of comments across categories of feedback use.

**Figure 3.**
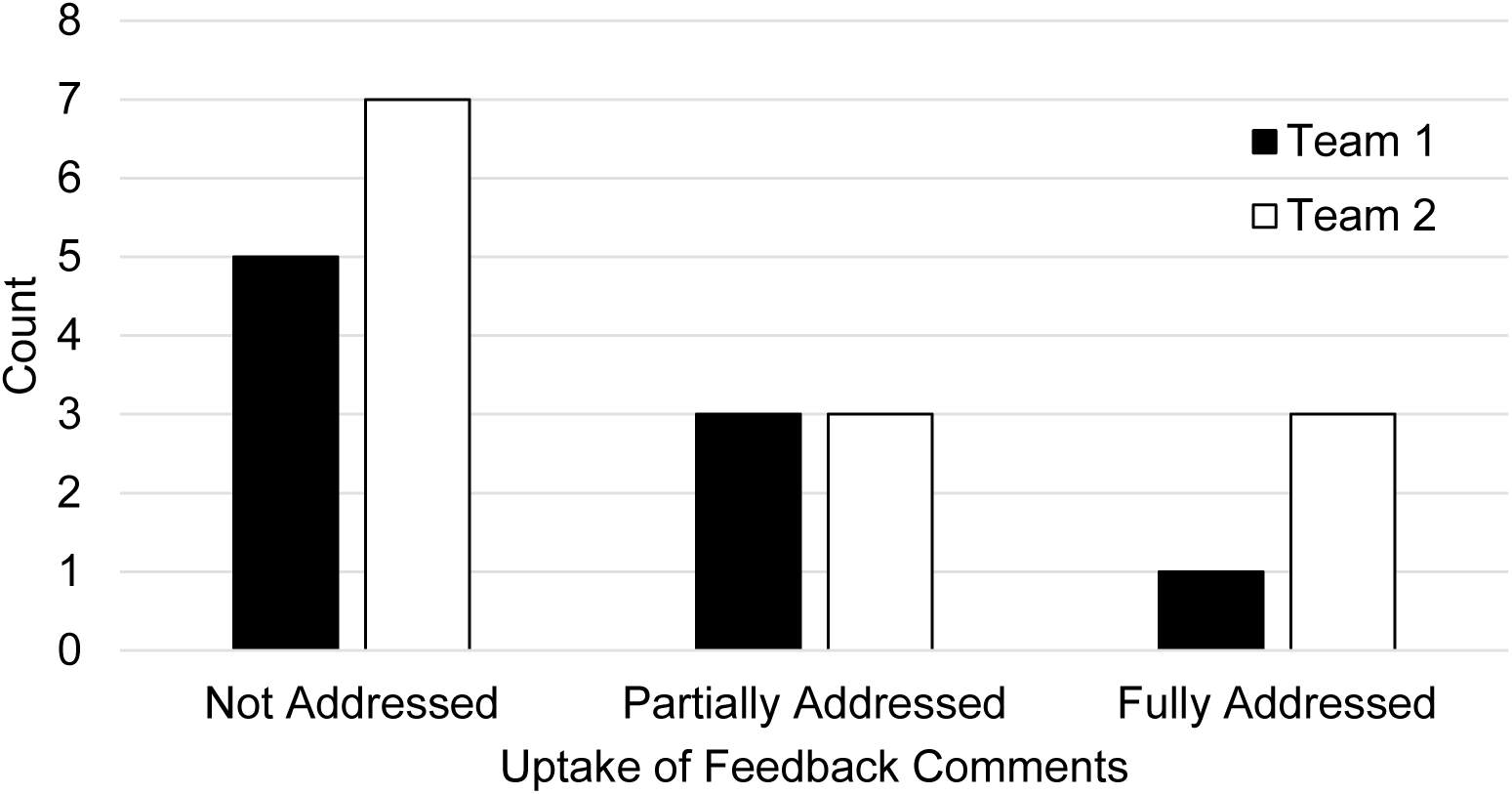
Uptake of feedback comments by student team.

### Team 1 Within-Case Analysis

Team 1 investigated the question, “In previously redlined communities, what is the relationship between tree density and median income, asthma frequency, and UV exposure?” In addition to examining the effects of redlining, their proposal introduction incorporated the construction of a nearby highway as a historically linked event with ongoing environmental and health consequences. They also described a local garden program as a community response initiative that could mitigate the harm perpetuated by redlining and the highway project. As noted earlier, Team 1 did not include a Micro paragraph in their first draft, and because the Micro paragraph in their final draft was newly written, it is excluded from this within-case analysis.

Overall, as the following analysis shows, Team 1 responded to AI-enabled feedback by improving conciseness and organization more than enhancing substantive content. Although feedback comments consistently requested additional information, explanation, reasoning, and evidence, Team 1 made only one addition to their final draft. This was a transition sentence that fully addressed the feedback comment suggesting they “…add a transition sentence between the Meso and Micro paragraphs to set up the shift from community impacts to individual outcomes.” In contrast, feedback comments that required adding new explanations or information largely went unaddressed. For instance, Team 1 did not address comments that asked them to 1) explain ways the local pattern of redlining compared to other cities, 2) explain how the community response initiative might relate to the study’s variables, or 3) better support claims with scientific reasoning or evidence, particularly those around the benefits of the community response initiative. Instead, the team most often responded to AI-enabled feedback through structural rather than conceptual revisions.

An additional example of how Team 1 avoided adding new information was by moving moving a large segment from the Meso Part 1 paragraph into the Macro paragraph. Collectively, three comments on the Macro paragraph had encouraged Team 1 to trace a timeline of consequences that would “help clarify *how* specific national practices led to systemic racial inequalities in housing and health that persist today”, beginning with the policies that institutionalized redlining. In response, they moved information about the highway project and its enduring impacts from the Meso Part 1 paragraph into the Macro paragraph. This shift partially addressed the feedback by extending the list of consequences. However, it did not conceptually connect the highway project to redlining nor elaborate on the policies that institutionalized redlining.

In other cases, Team 1 responded to feedback by removing text. For example, feedback advised that “Some claims—particularly around individual health outcomes and the benefits of the Garden Program—could be better supported with scientific reasoning or specific evidence”. The students responded to this comment indirectly by deleting an unsupported claim about the garden program in the Meso Part 2 paragraph. While this removal can be understood as an attempt to resolve the feedback by avoiding a weak claim, it did not involve adding reasoning or evidence. As such, the feedback was categorized as unaddressed.

Team 1’s revision pattern also reflected many deletions aimed at tightening the writing rather than expanding the content as comments had encouraged. In fact, Team 1 made so many deletions that the word count shrunk from 465 words in the first draft to 292 words in the final draft. As an example, the Macro paragraph originally contained two redundant explanations of redlining. The team removed the first explanation by deleting three text segments, leaving behind a sufficient introduction to redlining. This pattern extended across the proposal, as other deletions cut out unnecessary or redundant details, improving clarity and conciseness. In contrast, two of the team’s deletions of text from the Meso Part 1 paragraph removed details that were important for meeting rubric requirements. The rubric asked students to describe events that led to the perpetuation and exacerbation of inequities in ecosystems and human health. However, the team removed an explanation that the area’s historically lower tree density had resulted from redlining and disinvestment, and that economic barriers compounded to make it difficult for residents to mitigate harmful environmental effects. Similarly, the team deleted 5 citations that were essential for supporting claims. These examples show how the Team’s selective uptake of feedback is complex and involved competing goals for the revisions.

### Team 2 Within-Case Analysis

Team 2 chose to investigate the question, “How has redlining affected community garden quality and prevalence in the greater [city] area and how is it tied to obesity?” Their proposal focused on redlining and positioned obesity as a long-term health outcome tied to it. To unpack the relationship, they examined how poor soil quality and limited access to grocery stores in historically redlined areas contribute to elevated obesity rates. The team also incorporated community gardens that donate produce to a local food bank as a response initiative aimed at reducing environmental and health disparities.

The following analysis demonstrates that Team 2 responded to AI-enabled feedback by selectively expanding the proposal’s substantive content. One way Team 2 responded to AI-enabled feedback was by adding the explanations and details that comments requested. For example, feedback on the Meso paragraphs suggested they provide more description of the local food bank to clarify its role as a community response initiative, as the first draft had referenced it without supporting details. The team fully addressed the comments by adding a description that the food bank “aims to provide food pantries in the area with fresh produce. The [local food bank] is a non-profit organization that works with the community to provide food to those in need.” Team 2 also incorporated a small addition in their Macro paragraph after receiving feedback asking them to “explain how these policies caused long-term economic and health disparities. The connection between historical redlining and present-day diabetes is mentioned, but the causal chain could be more fully explained.” (Team 2 originally wrote about diabetes as another outcome variable before narrowing the scope to only obesity). They added the qualifier “limited” to describe healthcare access in the sentence, “This segregation has had lasting effects, contributing to health inequities like higher obesity and diabetes rates due to systemic barriers such as limited healthcare access and economic hardship.” Although their argument remained similar across drafts, the revision clarified the causal mechanism by specifying that healthcare access was less available in redlined areas.

In contrast to the addition of information requested by the feedback, Team 2 also made an addition that did not meaningfully address a feedback comment or help meet rubric criteria. In the Macro paragraph, they added the sentence, “Red and yellow areas, home to low-income Black residents, faced mortgage discrimination, while blue and green zones housed wealthier white populations.” While this elaboration identifies the color labels used in redlining maps, it does not explain how redlining’s short- and long-term consequences connect to present-day inequities, nor does it provide the context around the unique patterning of environmental and health conditions across neighborhoods in the specific city, as requested in another feedback comment. It is notable that the team incorporated this addition immediately after applying a scaffolding tip that provided a suggested sentence starter (“In the 1930s, redlining was codified through federal housing policy…”). Their revision reads, “In the 1930s, the Home Owners’ Loan Corporation (HOLC) and the Federal Housing Administration (FHA) institutionalized segregation through redlining, classifying neighborhoods into four color-coded zones. Red and yellow areas, …” A prior feedback comment also mentioned HOLC and FHA as historical entities, and it appears that the students found prescriptive or highly structured feedback easier to follow, whereas more open-ended feedback made it difficult for them to identify what substantive information to include next.

Another way Team 2 selectively responded to AI-enabled feedback was by focusing on paragraph-specific comments over broader suggestions aimed at improving the proposal’s flow and strength of arguments. Team 2 fully or partially addressed 6 of the 9 paragraph-specific comments, but none of the 4 whole-proposal comments. For example, a suggestion in the “Flow of writing” section noted that, “some paragraphs jump between topics (e.g., soil quality to obesity to garden location) without clear transitions. Strengthen the flow by using topic sentences that guide the reader and connect ideas more smoothly.” None of the team’s revisions improved transitions, so this feedback was categorized as unaddressed. Similarly, the “Claims are supported” section advised strengthening several claims with citations or scientific explanation and identified an example from the first draft that linked worms, soil quality, and hunger. Because none of the final draft’s added content provided evidence or reasoning—and because four citations from the first draft were deleted—these comments also remained unaddressed.

### Cross-Case Analysis

A comparison of the two within-case analyses shows similarities and differences in how students used AI-enabled feedback to revise their proposal introductions. The teams’ responses indicate that AI-enabled feedback can prompt different types of revision work: Team 1 primarily used it to refine structure through text deletions and moves, while Team 2 used it to elaborate content via additions. Despite these differences in revision strategies, the distribution of feedback comments across the categories fully, partially, and not addressed, was similar for both teams, suggesting comparable levels of uptake. This similarity in uptake reflects neither team consistently incorporating the conceptual information or reasoning that the feedback most often requested. Although Team 1’s structural revisions occasionally aligned with feedback—such as by moving information to a location that needed elaboration or removing unsupported claims—these changes did not expand the substantive content the feedback sought. Similarly, while Team 2 prioritized additions, they left many open-ended or conceptually demanding suggestions unaddressed. The remainder of the cross-case analysis describes the types of feedback that had limited use by both teams and offers possible explanations for the patterns observed.

A key source of limited uptake for both teams involved feedback that required identifying new information or conducting additional research. This pattern is shown on the vertical axis of Figure 4 where comments requiring common knowledge are more likely to be addressed than those that require information that must be researched. Across cases, comments that encouraged students to add location-specific historical details or to provide conceptual connections, scientific reasoning, or evidence were typically only partially addressed or left unaddressed. This pattern was especially apparent for Team 1, whose only addition was a transition sentence responding to a relatively prescriptive suggestion. Team 2 added new details—such as a description of the local food bank and the qualifier “limited” in their explanation of healthcare access— but these additions involved general knowledge that may not have required searching for external sources. For both teams, revisions that partially addressed feedback tended to meet the surface-level or easily actionable portion of a comment while omitting the more demanding element that required new evidence or conceptual elaboration. For example, Team 1 moved their description of the highway project into the Macro paragraph, which extended the list of consequences but did not include the causal explanation requested in the feedback (e.g., linking the highway project to redlining or describing the policies that institutionalized it). Incorporating new information into the proposal would have required multiple steps such as finding appropriate sources, assessing their relevance and quality, and integrating and citing them, and neither team demonstrated this type of engagement with the feedback. We interpret these missing or incomplete revisions as evidence that students opted to work with existing material over conducting additional research, suggesting either difficulty researching requested information or a reluctance to do so.

**Figure 4.**
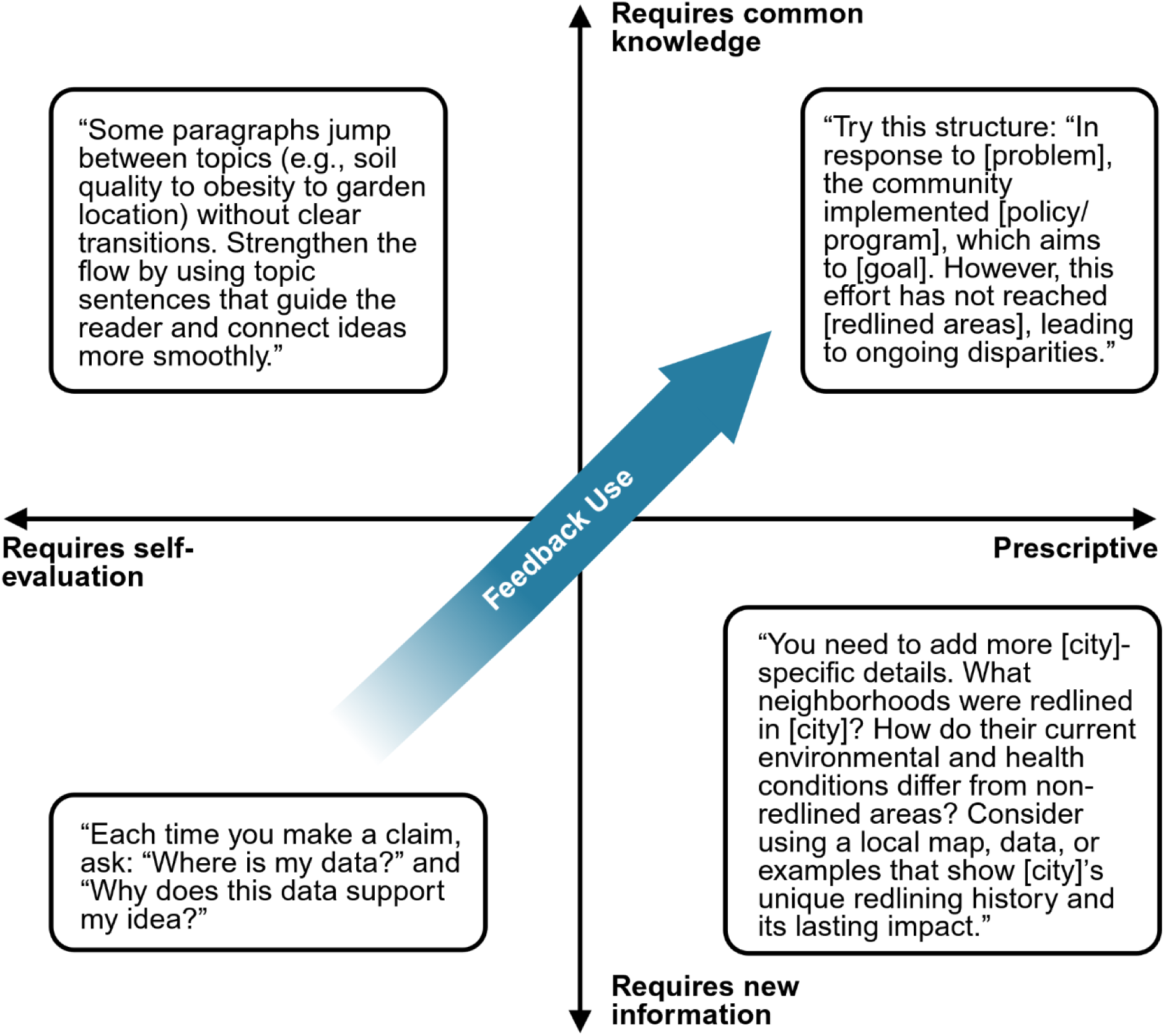
Factors Influencing Students’ Use of AI Enabled-Feedback

Both teams also made limited use of feedback that required self-evaluation, whether judging the coherence of the proposal, evaluating claims, or recognizing gaps in reasoning. This pattern is shown on the horizontal axis of Figure 4, where prescriptive comments are more likely to be addressed than those requiring self-evaluation. Except for Team 1’s new transition sentence, neither team addressed the feedback in the ‘Flow of Writing’ or ‘Claims Are Supported’ sections, which applied to the whole proposal. This suggests difficulty responding to whole-proposal comments, although the assignment’s design may have contributed; each student was responsible for writing one or two paragraphs, and they may have focused only on the feedback corresponding to paragraphs they authored. However, each whole-proposal suggestion included examples tied to specific paragraphs, and students still did not revise accordingly. The scaffolding tips in the ‘Claims are Supported’ section further illustrate the self-evaluative work expected of students. These tips directed them to examine their claims across the proposal and ask questions such as, “What’s the evidence?” and “Why does the evidence support the claim?” Neither team added support for claims, suggesting difficulty evaluating the adequacy of their reasoning in addition to the reluctance to research new information described previously. This difficulty identifying weaknesses in their explanations extended beyond the ‘Claims Are Supported’ section of feedback. As described in the previous paragraph, both teams often addressed only the easily actionable portions of feedback while neglecting the more conceptually demanding aspects, such as explaining the causal chain linking redlining to present-day inequities in the environment and human health.

Collectively, we interpret the observed patterns of feedback use as suggesting that students struggled not only to seek out new information but also to recognize conceptual gaps in their arguments, resulting in unsuccessful responses to some of the AI-enabled feedback. This interpretation is consistent with research showing that some students struggle to use feedback because they lack strategies for implementing it (Hyland, 1998; Winstone, Nash, Rowntree, et al., 2017). However, an alternative explanation is that students may use an informal cost-benefit analysis when deciding how much effort to invest in responding to different feedback comments. As a result, they may be less willing to devote the effort required to address more demanding feedback comments and make substantive revisions (Sutton, 2012; Winstone, Nash, Parker, et al., 2017). Notably, self-evaluation and finding new information typically require more effort than responding to prescriptive comments or comments that do not require additional research. Thus, students’ uneven use of feedback may reflect a combination of difficulties implementing feedback and a reluctance to invest effort in more demanding revisions.

### Theoretical Interpretation

AI-enabled feedback appears to support students’ engagement with revision, thus altering some, but not all, of the processes involved in feedback uptake. This section first examines evidence of students’ engagement with AI-enabled feedback, then explores barriers to its use, and finally synthesizes these patterns to characterize how it shaped feedback uptake for the two teams.

One way in which AI-enabled feedback’s support is evident is in students’ engagement with meaning-based revisions. Our findings show that both teams made meaning-based revisions in response to feedback, addressing roughly half of the feedback comments. This shows that students frequently engaged in substantive work rather than limiting their changes to surface-level edits, and is notable given well-documented challenges students face in using feedback to make substantive changes (reviewed in Jonsson, 2013; Winstone, Nash, Parker, et al., 2017). The prevalence of meaning-based revisions suggests that many AI-enabled comments were specific and interpretable enough to be useful. This interpretation aligns with prior research showing that students are more likely to act on feedback when it is explicit and detailed, whereas vague comments or symbolic markings can inhibit understanding and use (Ferguson, 2011; Jørgensen et al., 2023; Sutton, 2012). In this context, the relatively high rate of uptake suggests that the specificity and clarity of AI-enabled feedback may have supported students in revising their scientific writing in substantive ways.

Although both student teams revised their proposal introductions, we observed patterns in the ways in which they struggled to implement feedback. Students had difficulty evaluating their own reasoning, coherence of writing, and support for claims, and did not consistently incorporate the additional information or reasoning requested by the feedback. These patterns suggest that students did not always have the strategies, disciplinary understanding, or willingness to invest effort needed to act on them. This interpretation aligns with prior research indicating that feedback uptake depends not only on comprehension, but also on students’ self-assessment skills and their familiarity with discipline-specific terminology and practices (Jonsson, 2013; Nicol & Macfarlane-Dick, 2006; Sadler, 2010; Sutton, 2012; Winstone, Nash, Parker, et al., 2017). For example, feedback prompting students to evaluate the support for their claims may be intelligible, yet they may not have developed the self-assessment skills needed to act on it. They may also lack proficiency with the claim, evidence, and reasoning framework for scientific argumentation utilized in the assignment, which could prevent them from making adequate revisions despite targeted feedback. In addition, we observed a consistent pattern of lower uptake for feedback requiring further research or elaboration, suggesting that students were less likely to act on comments that demanded sustained effort or initiative. This pattern is consistent with research highlighting the role of volition in feedback use, including students’ willingness to invest effort and engage actively with feedback (Jørgensen et al., 2023; Winstone, Nash, Rowntree, et al., 2017). Across these patterns, we see that students often addressed the more easily actionable parts of AI-enabled feedback while omitting more demanding revisions that required self-evaluation or additional research and elaboration.

Taken together, these patterns suggest that the use of AI-enabled feedback supported students’ revisions primarily via the feedback’s content, while many of the challenges of implementing feedback remained. This tension underscores a learner-centered perspective of feedback, where students must actively interpret and implement feedback—not merely receive it—to benefit from it (Boud & Molloy, 2013). From this perspective, uptake depends on factors beyond the feedback itself, including students’ self-assessment skills, disciplinary understanding, and willingness to invest effort. Our findings suggest that these factors continue to constrain feedback use, indicating that several well-documented barriers to traditional feedback persist in this context. At the same time, patterns of uptake cannot fully account for all influences on students’ engagement with feedback. For instance, students’ use of AI-enabled feedback occurred within a broader context that included team dynamics, course workload, and individual motivation, each of which may shape how and whether feedback is taken up. This means that while AI-enabled feedback supported students in making meaningful revisions to their scientific writing, barriers persisted in its implementation, and likely in the broader context of the feedback process.

### Practical Implications

There are many barriers to providing formative feedback on scientific writing at scale, and to students’ use of feedback. In this section, we will describe practical implications of our findings for addressing some of the challenges faced by both students and instructors. Our analysis shows that both teams used some of their AI-enabled feedback, which is reassuring given how often students overlook or fail to read feedback (Hyland, 1998; MacDonald, 1991). This finding highlights the importance of providing opportunities for students to apply feedback through follow-up assignments like revised drafts. Providing students with an immediate opportunity to improve their work promotes their engagement with feedback (Paulus, 1999; Zhao, 2010). In addition, our observations suggest the potential value of requiring students to give written explanations of how they used each feedback comment, or to explain their rationale for not addressing it. Making students’ decision-making processes visible in this way may help instructors identify barriers to feedback use and more effectively provide support in future writing assignments.

We found that the custom AI tool used in this study primarily supported students’ feedback recipience through the content of the feedback itself, while barriers to its use remained. This implies that in addition to providing quality formative feedback, we also need to support students in implementing it. One way to do so is by refocusing scientific writing activities toward the development of feedback literacy. Our observed patterns in students’ feedback use indicate that uptake depends on their ability to evaluate their own work, their disciplinary understanding, and their willingness to invest effort in revision. These factors align with the underpinning features of feedback literacy—appreciating feedback, making judgements, managing affect, and taking action—suggesting that students may benefit from targeted instruction in this area (Carless & Boud, 2018).

Instructors often lack the time and resources needed to develop customized formative feedback, resulting in sparse comments or an uneven distribution across students. In practice, instructors may attempt to mitigate time constraints by prioritizing research proposals that show the largest gap in meeting rubric criteria, copy and pasting generic comments targeting common issues, or relying on peer feedback. While these strategies help manage workload, they can limit both the consistency and utility of feedback across students. The custom AI tool used in this study, grounded in learning theory and integrated with course-level supports, offers one potential way to address these challenges by supporting the development of comprehensive, instructor-vetted feedback for all students. The tool generated three forms of feedback for every paragraph of each proposal—strengths, suggestions for improvement, and scaffolding tips. This structure ensured that each team, regardless of how closely their work met the goals of the assignment, received detailed and actionable comments on their writing. Although this study did not aim to compare the time required to produce AI-enabled feedback versus traditional feedback, and our analysis is limited to two cases, the findings point to the potential for this approach to scale personalized formative feedback. Importantly, this approach may reshape how feedback is allocated. By ensuring that all students receive a baseline level of detailed, actionable feedback, AI-enabled feedback supports more equal access to support. At the same time, it may create opportunities for instructors to more deliberately direct additional support toward students with greater needs.

## Conclusions

GenAI tools are increasingly being used to support instructors in developing personalized formative feedback on students’ scientific writing. Therefore, it is necessary to examine how established understandings of feedback recipience and effectiveness apply in these AI-mediated contexts. In this study, we sought to describe and interpret how two teams of undergraduate biology students used AI-enabled feedback to revise their research proposal introductions. Our findings showed that both teams made substantive revisions after receiving AI-enabled feedback, addressing approximately half of the feedback comments. At the same time, both teams were less likely to implement certain types of feedback. In particular, they were less likely to address comments that required evaluating the quality of their reasoning, the coherence of their writing, or the adequacy of support for their claims. They also did not consistently incorporate the additional information or reasoning requested in the feedback. Taken together, these patterns suggest that the specificity and clarity of AI-enabled feedback supported meaningful revision, while challenges continued to shape how students implemented the feedback. Although the degree of uptake observed in this study is encouraging, our results suggest that feedback recipience—including recipience of AI-enabled feedback—is shaped by numerous learner-, task-, and context-related factors. Accordingly, it is unlikely that a single application of generative AI within the feedback process will resolve all the challenges associated with feedback uptake. To better understand how generative AI can support students’ use of feedback, future research should examine how AI tools shape specific aspects of the feedback process for both instructors and students, as well as how these tools influence learning outcomes. Future studies could also more directly investigate students’ perceptions of and challenges with AI-enabled feedback through qualitative interviews and written or verbal reflections on their responses to feedback. Overall, this study extends current understandings of feedback into the context of AI-enabled feedback and provides behavioral evidence of how integrating a genAI tool into the feedback process impacted students’ revisions.

## Supporting information

Supplemental Materials Weston Long

## Notes

This material is based upon work supported by the National Science Foundation Graduate Research Fellowship Program under Grant 2235783, the SimBio Foundation grant program, and the Plant Biology Department of Michigan State University.

### Competing Interest Statement

The authors have declared no competing interest.

